# Out of the wild: lab-reared bumble bees lack environmental bacteria and fungi

**DOI:** 10.64898/2026.11.24.768966

**Authors:** Danielle Rutkowski, María I. Pozo, Rachel L. Vannette

## Abstract

Social bees, including bumble bees, host a community of microbes in their gut consisting of core bacterial taxa as well as non-core bacteria and fungi. Bumble bee research colonies from commercial suppliers are often utilized in research characterizing and examining the functions of bee microbiomes. However, commercial rearing conditions have the potential to shift the microbiome associated with bees, calling into question whether results obtained using commercial colonies are applicable to wild bees. Here, we examine differences between the microbiome of commercially-reared and wild caught bumble bees of three species (*Bombus impatiens, B. terrestris,* and *B. vosnesenskii*) using amplicon sequencing and qPCR to characterize bacterial and fungal communities within bee guts. We found that while all bee species hosted some microbes belonging to the core bumble bee gut community, microbial community composition and diversity varied between species and sources, with wild individuals lacking some groups of “core” bacteria and hosting a greater diversity of non-core bacterial and fungal taxa. This work highlights the presence and potential dominance of non-core microbial taxa within wild bumble bee gut communities, especially early in the colony life cycle. Overall, we conclude that rearing conditions can shape the composition of bumble bee microbiomes, and research on commercial colonies alone may overlook environmental taxa commonly associated with bees.

## Introduction

For at least 4500 years, humans have managed bee colonies and populations for the products and services that they provide (Crane 1999). Today, several bee groups are managed for human use, including honey bees (*Apis*), bumble bees (*Bombus*), stingless bees (Meliponini), leafcutter bees (*Megachile*), mason bees (*Osmia*), and alkali bees (*Nomia*), all with varying degrees of human control and environmental exposure. Bumble bees are reared for pollination services of crops that are inefficiently or insufficiently pollinated by honey bees, including solanaceous crops like peppers, tomatoes, and eggplants, and ericaceous crops like blueberries and cranberries (Desjardins and Oliveira 2006, Morandin et al. 2001, Serrano & Guerra-Sanz 2006). Commercial bumble bee rearing operations began with the European species *Bombus terrestris* in 1987, and seven species (*B. atratus, B. canariensis, B. huntii, B. ignitus, B. impatiens, B. terrestris,* and *B. vosnesenskii*) are sold around the world today for crop pollination (Velthuis & van Doorn 2006, Koppert 2025). Unlike other managed bee species, commercially-produced bumble bee colonies are reared entirely indoors for the first 8-10 weeks of development, with no access to external foraging until they are installed in the field or greenhouse (Huang et al. 2016).

In addition to their pollination uses, managed colonies are also often used by researchers investigating questions in bee biology. These colonies are relatively inexpensive and can be procured in large quantities, making them preferable to wild or lab-reared colonies in many cases, as wild colonies are difficult to locate and wild-caught queens are time- and resource-intensive to rear with variable rates of successful establishment that can be as low as 0-5% for many species (Strange et al. 2023). Commercial colonies have been used in many experiments examining the structure and function of the bumble bee gut microbiome (Hammer et al. 2023, Palmer-Young et al. 2019, Nelson et al. 2025, Rothman et al. 2019, Rothman et al. 2020, Rutkowski et al. 2022, Straw et al. 2023). This microbiome, composed of a core set of bacterial taxa as well as non-core environmental bacteria and fungi, provides a number of benefits to the bumble bee host, including protection from pathogens and detoxification of toxic compounds (Koch & Schmid-Hempel 2011, Rothman et al. 2019). In honey bees, the microbiome performs several additional functions that are less well studied in bumble bees, including aided digestion of complex carbohydrates and stimulation of bee immune responses (Emery et al. 2017, Lee et al. 2015, Zheng et al. 2019).

As research on this topic continues to grow, it is important to examine the extent to which results obtained using commercially-produced bumble bee colonies are applicable to wild bee colonies. Specifically, we suspect that rearing and management conditions of commercial operations may alter the structure of associated microbial communities, likely due to a variety of factors. First, bumble bees from different sources are likely exposed to distinct microbial communities that differ in composition and diversity. Wild bumble bees are generalists and forage on a diversity of floral species to collect pollen and nectar, and these resources host microbial communities which in turn influence the microbial communities of bee nests (McFrederick & Rehan 2018, Rothman et al. 2019). In contrast, commercially-reared bees do not forage externally, and are fed artificial sugar solution and honey bee-collected pollen (Huang et al. 2016). While the sugar solution contains preservatives to inhibit microbial growth, certain osmophilic, preservative-tolerant microbes like the yeast *Zygosaccharomyces* are able to colonize this provision (Pozo et al. *in press*). In addition, honey bee-collected pollen is known to host a variety of microbes, including plant-associated microbes as well as bee parasites. These include *Crithidia spp*., *Apicystis bombi, Ascosphaera apis, Vairimorpha spp.,* and a variety of honey bee-associated viruses (Graystock et al. 2016, Pereira et al. 2019). Many of these parasites are known to infect bumble bees, leading to deformities, behavioral changes, disrupted colony founding and colony growth, or death in combination with other stressors (Gegear et al. 2005, Genersch et al. 2006, Maxfield-Taylor et al. 2015, Rutrecht & Brown 2008, van der Steen 2008). While some companies sterilize this pollen using gamma irradiation (Meeus et al. 2014), others use cycles of freezing and thawing that may be effective in reducing the abundance and viability of some microbes (Huang et al. 2016), though this method likely does not completely sterilize the provision (Pozo, unpublished results). This has led to concerns of disease transfer to wild, native pollinators, as commercially-reared bees may escape into the environment, acting as parasite vectors (Colla et al. 2006, Murray et al. 2013, Otterstatter & Thomson 2008). Honey bee-collected pollen may also contain pesticides (Mullin et al. 2010), and repeated exposure to this pollen may therefore apply different selection pressures in commercial rearing environments, possibly leading to microbiome divergence over generations. As the core bacterial microbiome of bumble bees is vertically transmitted between generations (Koch & Schmid-Hempel 2011), these aforementioned factors as well as limited inputs of new bee genetic material into rearing operations and variation in queen overwintering practices could potentially lead to altered gut microbiomes. Indeed, previous work suggests that the microbiomes of wild and commercial bumble bees therefore can differ (Hotchkiss et al. 2025, Meeus et al. 2015, Newbold et al. 2015), yet whether this difference is consistent among commercially reared species, is observed in queens (in addition to workers), and also applies to fungal communities remain unknown.

To determine differences between the microbiomes of commercially-reared and wild bumble bees, we characterized the gut-associated bacterial and fungal communities of three bumble bee species (*B. impatiens, B. terrestris,* and *B. vosnesenskii*). We used Illumina MiSeq amplicon sequencing to determine community composition and qPCR to determine total microbial abundance. We then tested for differences in community composition, diversity, bacterial and fungal abundance, and examined which taxa were differentially abundant between groups. We hypothesized: 1) the microbiomes of commercially-reared bees would be dominated by core gut bacteria, and would be less diverse than microbiomes of wild-caught bees; and 2) wild-caught bees would host a greater diversity of fungi and more variable fungal communities than commercially-reared bees.

## Methods

### Study species

Three bumble bee species (*Bombus impatiens, B. terrestris,* and *B. vosnesenskii*) were chosen for inclusion in this study due to availability of commercially-produced colonies and wild individuals. *B. impatiens* is a widespread and common species in the eastern United States, where it occurs in forest edges, fields, and meadows, including urban parks (Thuma et al. 2025, Williams et al. 2014). *B. vosnesenskii* is native to the western coast of North America, where it can be found in habitats spanning a wide elevation gradient, including chaparral and mountain meadows (Williams et al. 2014). *B. terrestris* is native to Europe, with a large Mediterranean-centered range (Rasmont et al. 2008), occurring in fields and woodlands (O’Connor et al. 2017, Svensson et al. 2000). In all three species, queens emerge from diapause early in the spring to found new colonies and generally nest underground in old rodent burrows (Plath 1922, Pugesek & Crone 2021, Thorp et al. 1983). All three species are generalists, foraging on a wide range of plant species (Cole et al. 2020, Kleijn & Raemakers 2008, Novotny et al. 2023). The history of commercial rearing operations varies by species, with *B. terrestris* sales beginning in 1987, *B. impatiens* in 1990, and *B. vosnesenskii* in 2021 (Velthuis & van Doorn 2006).

### Sample Collection and Processing

For each species, wild-caught and commercially-reared individuals were collected for comparison (Table 1). For *B. impatiens* and *B. vosnesenskii*, queens were collected for comparison, while workers were collected for *B. terrestris*. Wild *B. impatiens* queens were collected in Amherst, MA, USA in the spring of 2021 (n = 9). Commercial *B. impatiens* queens were obtained from a mixture of Koppert and BioBest colonies from April 2021 to September 2021 (n = 9). For *B. vosnesenskii,* wild queens were collected in March 2021 from Monterey, CA (n = 11), while commercial queens were obtained from Koppert colonies during this same time period (n = 5). *B. terrestris* workers were collected near Leuven, Belgium in May 2022 (12 wild, 12 commercial).

**Table 1.** Sampling locations and information for the three bumble bee species tested in this study. Wild queens were caught early in the season after emergence from hibernation, while commercial queens were sampled from fully established colonies.

| **Sampling location** | **Bee species** | **Bee source** | **Bee Caste** |
|---|---|---|---|
| Davis, CA, USA | *B. impatiens* | Commercial (Biobest, Koppert) | Queen |
| Winters, CA, USA | *B. vosnesenskii* | Commercial (Biobest) | Queen |
| Monterey, CA, USA | *B. vosnesenskii* | Wild | Queen |
| Amherst, MA, USA | *B. impatiens* | Wild | Queen |
| Leuven, Belgium | *B. terrestris* | Commercial (Biobest) | Worker |
| Schavotstraat, Begijnendijk, Belgium | *B. terrestris* | Wild | Worker |
| Papendel, Begijnendijk, Belgium | *B. terrestris* | Wild | Worker |
| Betekom, Belgium | *B. terrestris* | Wild | Worker |

Following collection, all bees were frozen at -80°C until dissection. The midgut and hindgut were dissected from each bee, excluding the crop due to previously observed low microbial density and risk of PCR inhibition. DNA was extracted from samples using a QIAGEN DNeasy PowerSoil Pro DNA extraction kit (for *B. impatiens* and *B. vosnesenskii*) or Norgen Biotek Plant/Fungi DNA isolation kit (for *B. terrestris*), following included kit instructions. All DNA was quantified on a NanoDrop One (Thermo Scientific, USA) to ensure sufficient quantity and quality, and was stored at -80°C prior to sequencing.

### DNA Sequencing and Quantification

Samples were sent to the Integrated Microbiome Resource at Dalhousie University for Illumina MiSeq amplicon sequencing of fungal and bacterial communities. For fungal communities, the ITS2 region was sequenced using the ITS86F (5’ GTGAATCATCGAATCTTTGAA 3’) and ITS4R (5’ TCCTCCGCTTATTGATATGC 3’) primers (Op De Beeck et al. 2014). For bacteria, the V5-V6 16S region was sequenced using the 799F-mod3 (5’ CMGGATTAGATACCCKGG 3’) and 1115R (5’ AGGGTTGCGCTCGTTG 3’) primers (Hanshew et al. 2013). To determine microbial abundance, qPCR was run on all samples using a CFX96 Touch Real-Time PCR Detection System to determine total 18S (for fungi) and 16S (for bacteria) gene copy number. For bacteria, the same primers were used as above, with the following cycle conditions: 3 min at 95°C followed by 35 cycles of 30s at 95°C, 30s at 52°C, and 1 min at 72°C. For fungi, the fungi-specific Taqman protocol FungiQuant (Liu et al. 2012) was used to quantify 18S gene copies using the following cycle conditions: 10 min at 95°C followed by 45 cycles of 15s at 95°C and 1 min at 65°C. To convert from Cq values obtained from qPCR to total copy number, plasmids with 16S (*Nocardioides luteus*) and 18S (*Moniliella oedocephalis*) gene sequence inserts were used (Eurofins Genomics Blue Heron). Standard curves were created using dilution series of the 16S plasmid (R^2^ = 0.98) and the 18S plasmid (R^2^ = 0.98), and were used to convert sample Cq to total copy number.

Illumina ITS and 16S sequence data were processed using the dada2 pipeline in R (package ‘dada2’, Callahan et al. 2016). For 16S sequences, the default dada2 pipeline parameters were followed and taxonomic assignments were made using the BEExact database (v2023.01.30, Daisley & Reid 2020). Due to poor assignments of common contaminants (such as chloroplast or mitochondria DNA) using this database, the Silva database (v138.1, Chuvochina et al. 2026) was also used for taxonomic assignment, and results were compared. For mismatched assignments, sequences were identified using nucleotide BLAST using the standard core nucleotide database. Top BLAST assignments were identified using E-values and percent identity. These assignments were then used in further processing. For all other ASVs, the BEExact assignment was used (Table S1). For ITS sequences, cutadapt (Martin 2011) was used to remove primers from sequences before further processing. Default parameters were used throughout the pipeline with the exception of the merging step, for which one error was allowed between forward and reverse sequence matches. Taxonomic assignments were made using the UNITE database (version 29.11.2022, Nilsson et al. 2018). Sequences obtained from Illumina sequencing were filtered to remove non-target reads prior to further analysis. Contaminants were removed from samples using the prevalence method in the ‘decontam’ package (Davis et al. 2017). From 58 bee gut samples and following filtering, we recovered a total of 1,707,517 16S reads, with a mean of 29,440 reads per sample (range 826 -66,790 reads). These clustered into 1022 total ASVs, with a mean of 28 ASVs per sample (range 3 -251 ASVs). We recovered 232,309 ITS reads following filtering, with a mean of 4,005 reads per sample (range 0 - 22,179 reads). The number of ITS reads recovered was much higher in wild *B. terrestris* compared to all other sample types (species * source interaction, F_2,55_ = 16.7, p < 0.001). Samples with fewer than 500 reads were filtered from the dataset, resulting in the loss of all but 1 commercial *B. vosensenskii* sample and 2 commercial *B. impatiens* samples. Because of this low sample number, all subsequent analyses testing the effect of bee source on fungal community were run using *B. terrestris* data only. Recovered ITS reads clustered into 572 ASVs, with a mean of 33 ASVs per sample (range 1 - 79).

### Statistical Analysis

All data analysis was performed in R version 4.2.2 (R Core Team 2022). Unless otherwise specified, bee species, source, and their interaction were included as predictors in all analyses. The proportion of reads belonging to core genera were calculated for each sample and compared using ANOVA. For these analyses, core bacteria were defined as those belonging to the genera *Gilliamella, Snodgrassella, Schmidhempelia, Lactobacillus, Bombilactobacillus, Apilactobacillus, Bifidobacterium,* and *Bombiscardovia* (Kwong et al. 2017). To determine how bacterial and fungal abundance differed across bee species and source, log-transformed 16S and 18S copy numbers were analyzed using ANOVA. Using the ‘phyloseq’ (McMurdie & Holmes 2013) and ‘microbiome’ (Lahti et al. 2017) packages, alpha diversity metrics (observed richness, Shannon diversity, Pielou’s evenness) were calculated and analyzed using ANOVA. For diversity analyses, known core gut bacteria were analyzed separately from non-core taxa.

Differences in microbial community composition were analyzed using PERMANOVA on calculated Bray-Curtis dissimilarities, using 999 permutations. Differences in community variance were tested using betadisper tests in the ‘vegan’ package (Oksanen et al. 2025). For analyses of fungal community composition and variance, separate analyses were run to test the effects of bee species and bee source, and only *B. terrestris* samples were used for comparisons between sources. Differentially abundant bacterial and fungal taxa between commercially reared and wild caught bees were determined using differential gene expression analysis in package ‘DESeq2’ (Love et al. 2014). Differential expression analysis was performed separately for each species, with bee source as a predictor. For determining differentially abundant fungal taxa between commercial and wild sources, only *B. terrestris* samples were analyzed. For significant results, posthoc analyses were carried out between treatment groups using estimated marginal means (package ‘emmeans’, Lenth & Piaskowski 2025)

## Results

### Wild bees host greater diversity and relative abundance of non-core bacteria

Across bee species, bacterial communities were generally dominated by core taxa, which comprised 79% of total reads, though this varied greatly between bee species (F_2,52_ = 14.1, p < 0.001) and bee source (F_1,52_ = 25.7, p < 0.001, Figure 1), with particularly low proportions of core reads in wild *B. vosnesenskii* queens. The top genera across bee species were *Snodgrassella* (22% of total reads), *Lactobacillus* (19%), and *Schmidhempelia* (14%). Bacterial community composition was significantly impacted by bee species (F_2,52_ = 9.11, R^2^ = 0.21, p = 0.001), source (F_1,52_ = 5.61, R^2^ = 0.066, p = 0.001), and their interaction (F_2,52_ = 4.93, R^2^ = 0.11, p = 0.001, Figure 2, Figure S1A). Commercial *B. impatiens* queens hosted more diverse communities of core gut bacteria than their wild counterparts (Figure 1, Figure S3) but did not differ in overall bacterial diversity (Figure S2). Wild *B. impatiens* queens hosted greater diversity and relative abundance of non-core bacteria, as well as more even communities (Figure 1, Figure S3). For *B. vosnesenskii* and *B. terrestris*, limited effects of bee source on Shannon diversity were observed, though the richness of core bacterial communities varied by species and source (Figure S3). Community variance was unaffected by bee species (F_2,55_ = 0.28, p = 0.75) and was only marginally impacted by bee source (F_1,56_ = 3.87, p = 0.055), with wild-caught bees hosting more variable bacterial communities.

**Figure 1.**
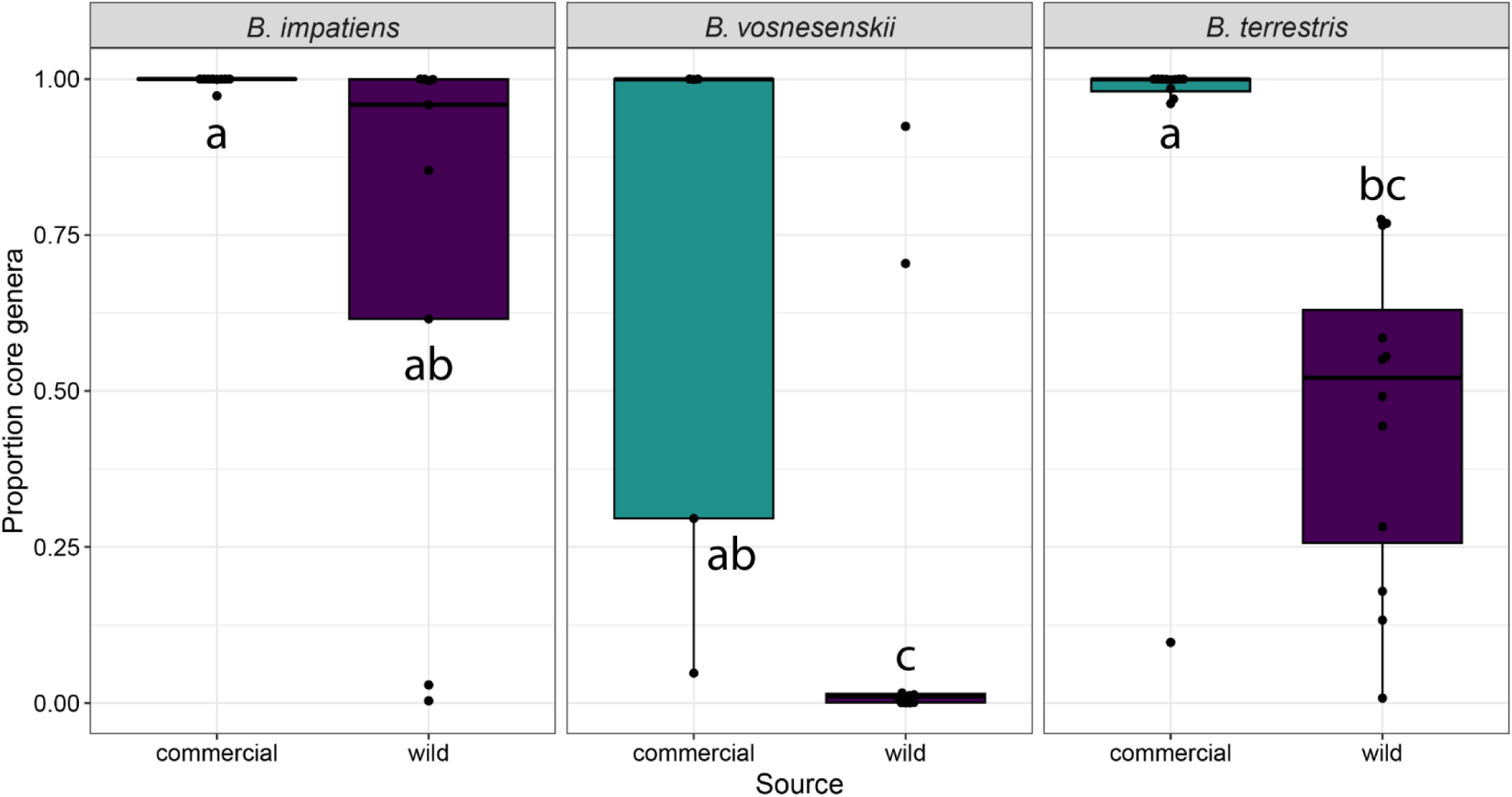
The proportion of 16S reads belonging to core bacterial genera across three bumble bee species from wild and commercial sources. Bee species (F_2,52_ = 14.1, p < 0.001) and source (F_1,52_ = 25.7, p < 0.001) impacted the prevalence of core reads across samples. Different letters represent statistically different values as determined by estimated marginal means (α = 0.05).

**Figure 2.**
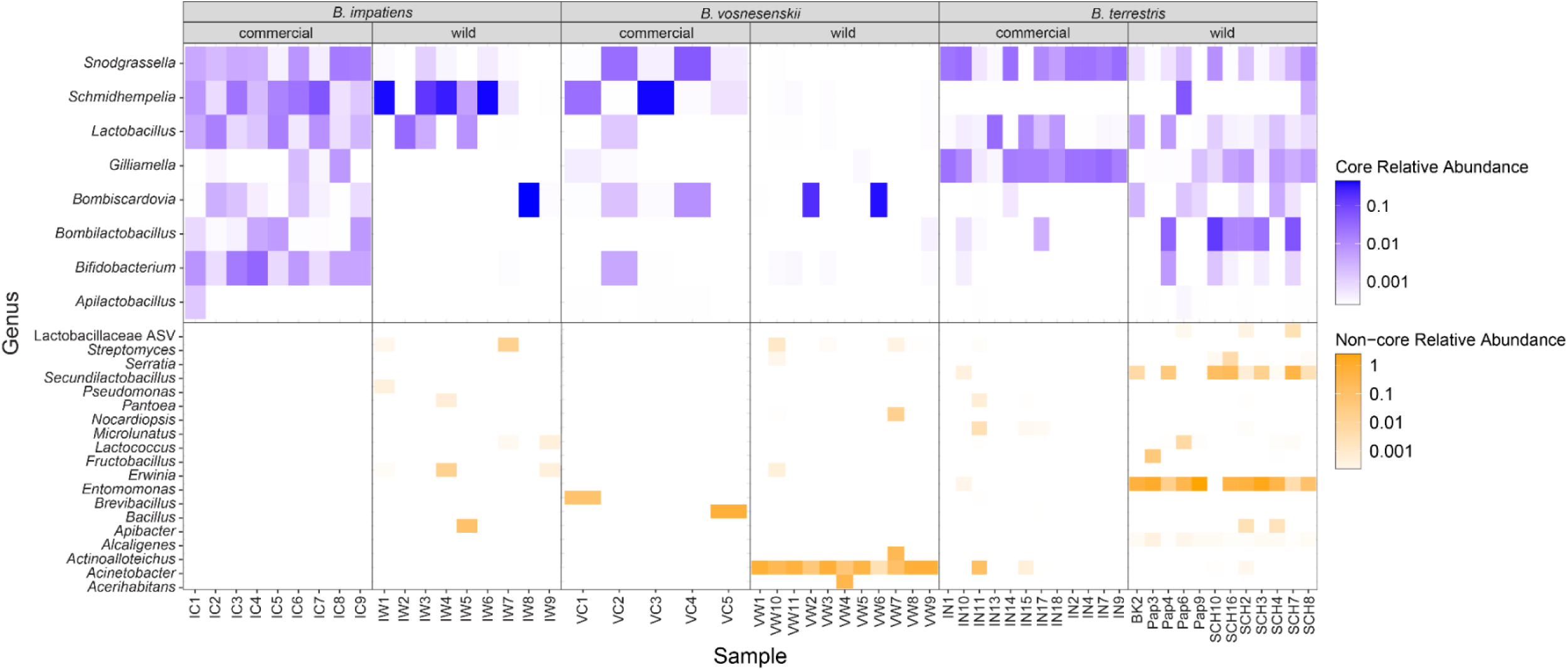
Bacterial community composition of three different bumble bee species from commercial and wild sources. For core genera, the summed abundance of all ASVs within each genus are shown, while the top 50 non-core ASVs are shown. Core genera are represented in blue, and non-core genera are represented in orange.

DESeq2 analysis revealed numerous differentially abundant taxa between commercial and wild bees. In most cases, ASVs of core genera were enriched in commercially-sourced bees compared to wild bees, while non-core genera were more abundant in wild bees (Figure S4). In *B. impatiens*, the genera *Bifidobacterium, Bombilactobacillus, Bombiscardovia, Gilliamella,* and *Lactobacillus* were more abundant in commercial bees, while *Erwinia, Streptococcus,* and *Streptomyces* were more abundant in wild bees. *Gilliamella*, *Bombilactobacillus* and *Apilactobacillus* were completely absent in wild *B. impatiens*. In *B. terrestris, Gilliamella* and *Microlunatus* were more abundant in commercial bees, while *Alcaligenes, Bifidobacterium, Entomomonas, Lactobacillus, Schmidhempelia,* and a Lactobacillaceae ASV were more abundant in wild bees. In *B. vosnesenskii*, *Bombiscardovia* and *Snodgrassella* were more abundant in commercial bees, and *Acinetobacter, Actinomycetospora, Bifidobacterium, Mycolicibacterium, Nakamurella,* and *Streptomyces* were more abundant in wild bees. *Gilliamella* and *Apilactobacillus* were completely absent in wild *B. vosnesenskii*. In wild individuals of *B. vosnesenskii* and *B. terrestris*, non-core bacteria comprised major portions of the microbiome, with *Acinetobacter* comprising 65.9% of reads on average in wild *B. vosnesenskii* queens, and *Entomomonas* comprising on average 37.6% of reads in wild *B. terrestris* workers.

### Wild bees often host greater fungal diversity and distinct species compared to commercial counterparts

ITS reads were dominated by fungi in the phyla Basidiomycota (66.9% of reads) and Ascomycota (32.8% of reads). Fungal community composition was extremely variable in *B. impatiens* and *B. vosnesenskii*, and normally dominated by just a few ASVs of *Aureobasidium, Cladosporium, Debaryomyces, Pseudogymnoascus, Schwanniomyces, Verrucocladosporium,* and *Zygosaccharomyces*. In *B. terrestris*, samples were much more consistent in their fungal composition, with commercial samples dominated completely by *Saccharomyces* while wild samples hosted consistent communities of *Luellia* and ASVs belonging to the Agaricomycetes and Russulales (Figure 3). Fungal ASV richness and Shannon diversity were higher in wild *B. terrestris* compared to their commercial counterparts (Figure 4). Bumble bee species also impacted fungal community diversity, with *B. vosnesenskii* hosting more diverse and more even communities (Figure S5).

**Figure 3.**
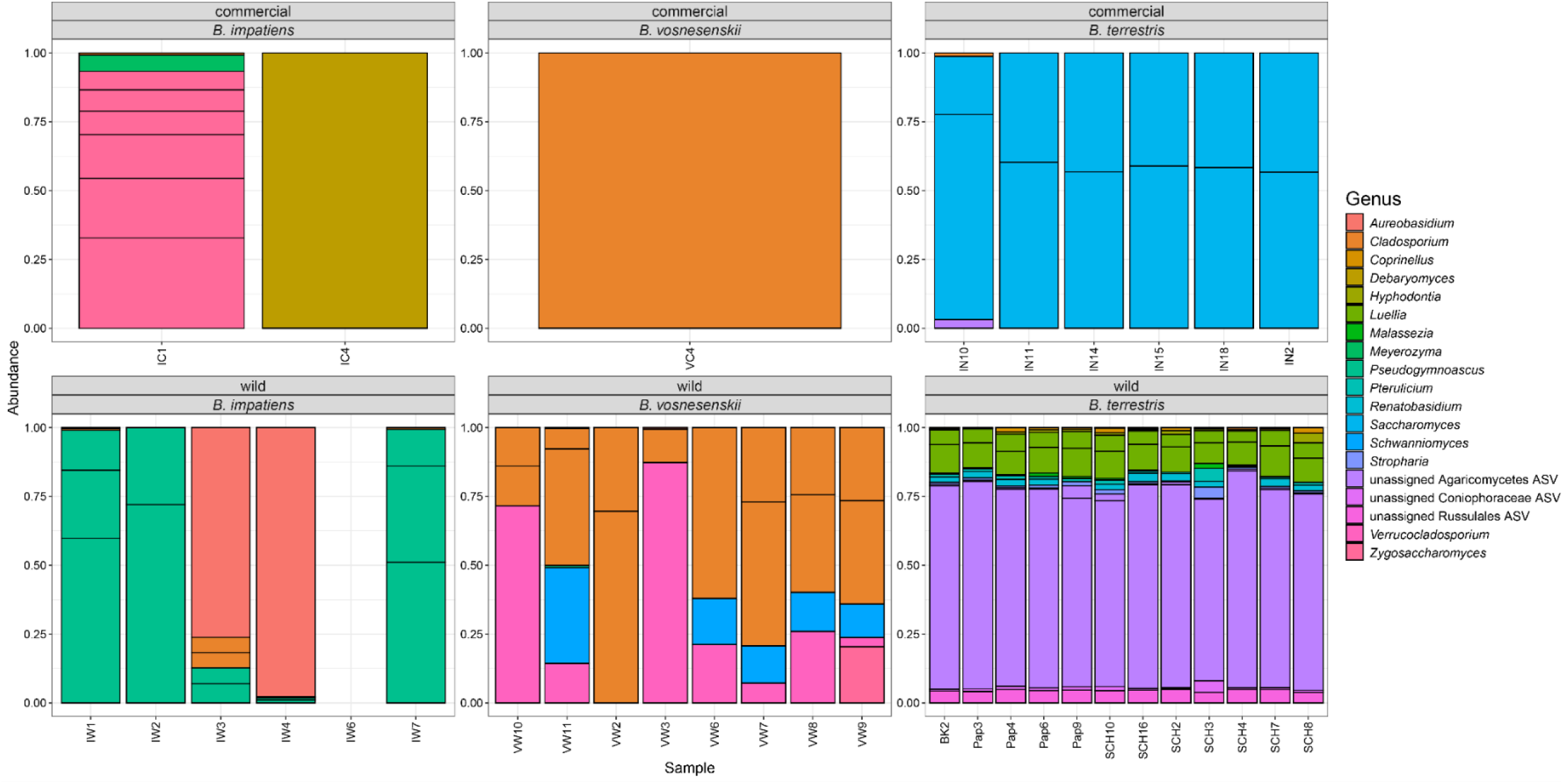
Fungal community composition of three different bumble bee species from commercial and wild sources. The top 30 ASVs are represented. Some samples are not represented, as they contained only less abundant taxa that were not captured in the top 30 ASVs.

**Figure 4.**
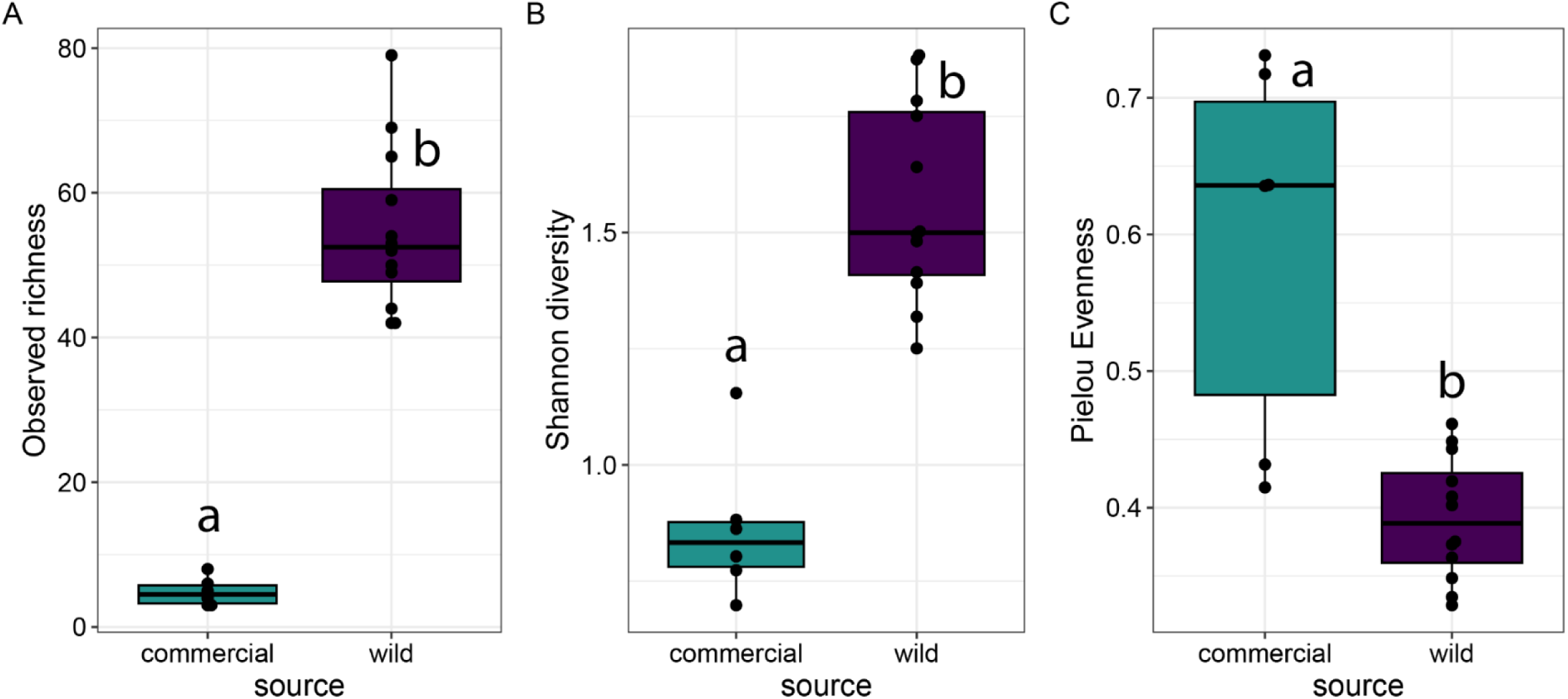
Observed richness (A), Shannon diversity (B), and Pielou’s evenness (C) of fungal gut communities of *B. terrestris* workers from commercial and wild sources. Bee source significantly impacted observed richness (F_1,16_ = 111, p < 0.001), Shannon diversity (F_1,16_ = 49.9, p < 0.001), and Pielou’s evenness (F_1,16_ = 22.1, p < 0.001) of fungal communities. Different letters represent statistically different values (α = 0.05).

Fungal community composition varied between commercial and wild workers of *B. terrestris* (F_1,16_ = 357, R^2^ = 0.96, p = 0.001, Figure 3, Figure S1B), while community variance was similar across bee sources (F_1,16_ = 0.0036, p = 0.94). Only two ASVs, both belonging to *Saccharomyces*, were more abundant in commercial bees, while wild bees were characterized by the presence of multiple ASVs, including ones belonging to Agaricomycetes, Russulales, and *Luellia* (Figure S6). Fungal communities also differed between bee species (Figure S7).

### Source effect on microbial abundance is limited for bacteria but strongly affects fungi in B. vosnesenskii

Bacterial abundance, measured using total 16S copy number, averaged 5.58 x 10^5^ 16S copies per sample and did not differ among bee species (F_2,52_ = 0.58, p = 0.56), bee source (F_1,52_ = 1.18, p = 0.28), or their interaction (F_2,52_ = 0.59, p = 0.56, Fig 5A). In contrast to bacteria, fungal abundance was strongly affected by source but only in *B. vosnesenskii*, with wild caught queens hosting 1,400 times higher 18S copies compared to commercially sourced queens (species * source interaction, F_2,52_ = 12.52, p < 0.001, Fig 5B).

**Figure 5.**
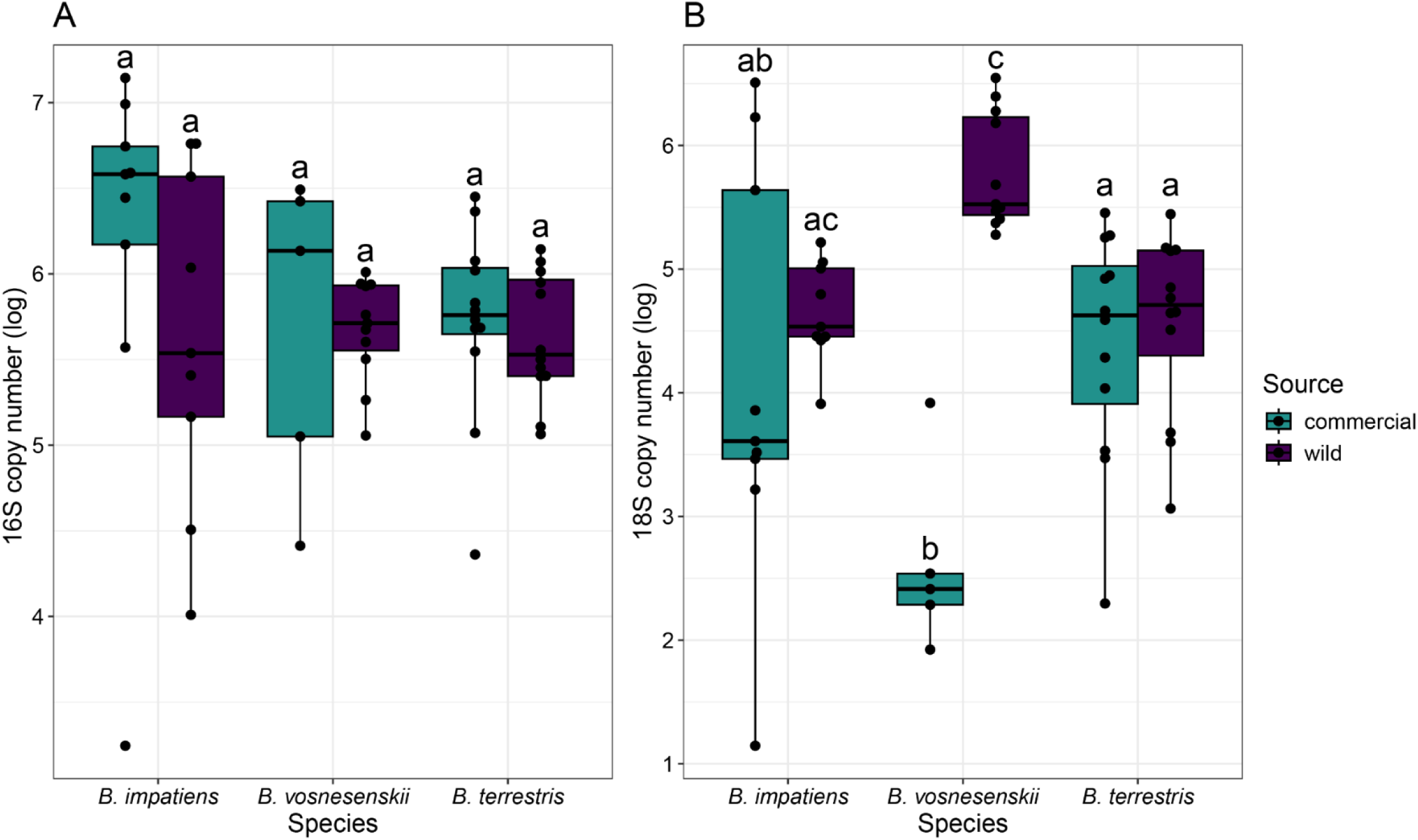
Total abundance of A) bacteria (16S copy number) and B) fungi (18S copy number) in three species of bumble bees from commercial and wild sources. 16S copy number was unaffected by species or source (p > 0.05 for all comparisons), while 18S copy number was impacted by the interaction between these two factors (F_2,52_ = 12.5, p < 0.001). Different letters represent statistically different values as determined by estimated marginal means (α = 0.05).

## Discussion

We characterized gut microbial communities of three bumble bee species from commercial colonies and wild sources to determine if commercial rearing impacts the bee microbiome. Despite significant overlap in core microbiome members, bacterial and fungal communities of bee guts varied between bee species and sources, with wild bees generally hosting a greater diversity of non-core bacteria and fungi. These differences between commercial and wild microbiomes were also dependent on bee species. In particular, we saw large differences between wild-caught, early season queens of *B. impatiens* and *B. vosnesenskii* compared to commercially reared queens from established colonies, which could be due to timing, rearing conditions, or both. For *B. terrestris* workers, wild bees hosted *Entomomonas* and a consistent basidiomycete fungal community, both of which were absent in commercially-sourced workers and are possibly sourced from nesting habitats.

Our findings generally agree with previous studies examining the role of commercial rearing and outside foraging on the microbiomes of bumble bees, which often find a higher diversity and abundance of environmental microbes in bees caught from the wild or given access to outside foraging (Meeus et al. 2015, Newbold et al. 2015, Weinhold et al. 2024). Commercial colonies of *B. terrestris* given access to environmental foraging host similar or even greater levels of microbial diversity than wild-caught conspecifics (Newbold et al. 2015) and when commercial colonies of *B. terrestris* were moved outdoors and allowed to forage, microbiomes shifted, with increased diversity over time (Weinhold et al. 2024) or a dramatic shift in composition (Parmentier et al. 2016). Additionally, wild-caught queens reared indoors on the same diet as commercial bees develop microbiomes that are dominated by core bacteria, with no environmental microbes present (Hammer et al. 2023), further supporting the importance of rearing conditions on microbiome structure. In contrast, some studies observed no differences in bacterial diversity between indoor- and outdoor-reared bumble bees (Hotchkiss et al. 2025, Parmentier et al. 2016, Roy et al. 2023). However, these studies looked at overall bacterial diversity, while in the current study we only found significant changes in diversity when analyzing core and non-core bacteria separately. Given that several of these studies did note compositional shifts between bee groups, it is possible that there were shifts in the diversity of core and non-core bacteria that were not captured. Additionally, differences between studies may in part be due to OTU filtering parameters, as some studies used stricter abundance-based cutoffs for OTU inclusion in analyses (Parmentier et al. 2016, Roy et al. 2023), which may exclude rarer environmental microbes present at lower relative abundances.

In the current study, we observed that wild individuals of *B. terrestris* and *B. vosnesenskii* hosted microbiomes composed of a high proportion of environmental bacteria. *Entomomonas* was detected in all but one wild *B. terrestris* sample at high relative abundance (37.6% ± 7.6% SE), and *Acinetobacter* dominated wild *B. vosnesenskii* queens, comprising 65.9% of 16S reads. *Entomomonas* has recently been recognized as a common associate of insects, found in association with diverse insect groups including bees, butterflies, flies, crickets, and beetles (Busse et al. 2021, Comeault et al. 2025, Moser et al. 2022, Wang et al. 2019, Weinhold et al. 2026). *Acinetobacter* bacteria are common associates of floral nectar and bees (Alvarez-Perez et al. 2021, Fridman et al. 2012, Sless et al. 2026, Tang et al. 2021), and may attract pollinators to flowers in concert with nectar yeasts (Colda et al. 2021). Other studies on *B. terrestris* and *B. impatiens* have identified similarly high levels of environmental bacteria taxa in wild or free-foraging workers, including *Arsenophonus*, *Fructobacillus*, and Enterobacteriaceae (Hotchkiss et al. 2025, Krams et al. 2022, Meeus et al. 2015, Nelson et al. 2025, Parmentier et al. 2016). These gut microbiome enterotypes dominated by environmental bacteria were identified in 25% of samples in a survey of 28 bumble bee species, suggesting that they may be relatively common in wild bees (Li et al. 2015). It is also possible that bee caste and timing of sampling impacted the recovered microbiomes in the current study, as wild queens of *B. impatiens* and *B. vosnesenskii* were sampled early in the season, when queens had just emerged from diapause to found new colonies. Other studies on the microbiome of diapausing *B. impatiens* and *B. terrestris* queens have found that gut microbiomes are disturbed during overwintering, with lower bacterial abundance (Hotchkiss et al. 2024) and reduction or loss of core microbiome members such as *Snodgrassella, Gilliamella,* and *Bifidobacterium* (Bosmans et al. 2018a, Hotchkiss et al. 2024, Koch et al. 2013). This is in line with what we observed in our wild-caught queens, which hosted lower abundance of core microbiome groups like *Bifidobacterium, Bombilactobacillus, Bombiscardovia, Gilliamella,* and *Snodgrassella* compared to their commercial counterparts, which may have allowed environmental bacteria like *Acinetobacter* to dominate. As we saw no effect of bee source on bacterial abundance in the gut, these differences seem to be due to a high abundance of *Acinetobacter* in these early queens, as opposed to a reduction in total microbiome abundance and therefore increased detection of less common microbes. However, it is worth noting that the average bacterial abundance that we recovered in this study (5.57 x 10^5^ 16S copies) is orders of magnitude lower than those found in previous studies of *B. impatiens* workers, which average close to 10^8^ 16S copies (Hammer et al. 2021). This disparity could be due to the use of different bumble bee species and castes in the current study, or it could be that the methods we used here for bacterial quantification underestimate abundance. Work on *B. lantschouensis* found that while mated queens were dominated by non-core bacteria, post-diapause queens sampled just after first worker emergence hosted more core-dominated microbiomes (Wang et al. 2019), suggesting that microbiomes disturbed through diapause may converge on a consistent microbiome. However, research on bee microbiomes that uses commercial colonies alone may underestimate the frequency and importance of environmental microbes, and more research would inform their impacts on colony success.

Fungal communities also differed between wild and commercial bees, particularly in *B. terrestris*. Fungal associations of *B. impatiens* and *B. vosnesenskii* were found to be sporadic and inconsistent, with most bee samples dominated by one or two fungal taxa, including *Cladosporium, Veruccocladosporium,* and *Psuedogymnoascus*. In *B. terrestris*, commercial workers were dominated by *Saccharomyces,* which may reflect their diet, as this yeast can be used as an additive to the diets of managed bees (Dodologlu & Emsen 2007, Fernandes-da-Silva & Zucoloto 1990, van der Steen 2007). Wild *B. terrestris* workers, however, hosted mostly Basidiomycetous fungi, including *Luellia* and an unidentified Agaricomycete fungus. These fungi are often decomposers of wood and are common in soils, and therefore may be associated with nesting sites of wild *B. terrestris*. Though these species have not been described in wild-caught queens of *B. terrestris* or other social bee species (Bosmans et al. 2018b, Brysch-Herzberg 2004, Pozo et al. 2018, Rutkowski et al. 2023), previous work found that only forest-caught *B. terrestris* queens contained detectable fungi, including soil fungi *Trichoderma* and *Mucor*, suggesting some fungal acquisition from environmental sources (Bosmans et al. 2018b). Our sampled bees were from sites approximately 2-2.7km apart, a distance which is at or beyond the maximum recorded foraging ranges of *B. terrestris* workers (Knight et al. 2005, Kreyer et al. 2004, Walther-Hellwig & Frankl 2003), making it unlikely that all our collected workers were from the same nest. This suggests that these fungal associations may be consistent across different nests. Other mechanisms of acquisition are also possible, as bees have been observed to directly collect fungal spores while foraging (Eltz et al. 2002, Shaw 1990), and addition of oyster mushroom mycelia to *B. impatiens* diets has been shown to improve colony development (Donzelli & Strange 2025), so more investigation into the consistency of this fungal association in *B. terrestris* and the impacts of these fungi on bee health are necessary.

In addition to differences between wild and commercial bees, we also observed differences in microbiome composition between bee species, as expected based on previous studies (Cariveau et al. 2014, Hotchkiss et al. 2025, Kwong et al. 2017, Villabona et al. 2023). Core bacterial taxa were very prevalent in both commercial and wild samples of *B. impatiens*, comprising 99.6% and 92.3% of 16S reads, respectively. This dominance of core taxa has been demonstrated in previous studies comparing commercial and wild *B. impatiens* (Hotchkiss et al. 2025). However, we observed different patterns in *B. terrestris* and *B. vosnesenskii* samples in this study. While commercial *B. terrestris* workers were also dominated by core bacteria (97% of 16S reads), these bacteria made up only 50% of reads in wild workers. The proportion of core bacteria reads was even lower in *B. vosnesenskii*, comprising 61% of reads in commercial queens and only 29% of reads in wild queens. Previous studies on *B. terrestris* show similar increases in the prevalence of non-core bacteria in wild individuals or workers given access to outside foraging (Meeus et al. 2015, Newbold et al. 2015, Parmentier et al. 2016), though some studies find a dominance of core bacteria regardless of bee source (Hotchkiss et al. 2025, Weinhold et al. 2024), suggesting that the microbial community of the environment as well as other environmental factors may play a role in determining this shift. The particularly low abundance of core microbes in commercial *B. vosnesenskii* may be due to the fairly recent commercialization of this species, with sales beginning in 2021 (Koppert, 2021). As wild queens are introduced into captive rearing operations during the first several years (Huang et al. 2016), they possibly also introduce microbiomes dominated by environmental bacteria. As queens are the main source of microbes for a developing colony (Su et al. 2021), and the core microbiome normally establishes in adult bumble bees within 4 days of eclosion (Hammer et al. 2023), it is possible that early exposure to certain environmental microbes encountered in the bee nest and surrounding environment, or other environmental factors, may disrupt normal microbiome assembly (Nelson et al. 2025).

Overall, we found differences in bacterial and fungal diversity and composition between bumble bees from different sources, with some environmental microbes dominating early season queens and foraging workers. These results suggest that environmental microbes may be important constituents of the microbiome in multiple portions of the bumble bee life cycle, and these microbes may be overlooked in studies performed exclusively on commercial bumble bee colonies. While the metabolic potential of core-dominated and non-core-dominated bumble bee microbiomes were found to be similar in a study of four bumble bee species (Hotchkiss et al. 2025), whether there are any other differences in microbiome function or impacts on bee hosts between these microbiome types remains to be tested.

## Author Contributions

DR and RLV conceived of the study. DR and MIR collected and processed samples. DR processed sequences, performed statistical analyses, and wrote the first manuscript draft. DR, RLV, and MIR contributed to revisions. All authors approved of the submission.

## Acknowledgements

We thank Lynn Adler for the contribution of bee samples to this study.

