## Supplemental Material for "Out of the wild: lab-reared bumble bees lack environmental bacteria and fungi"

Table S1. Discrepancy between Silva and BEExact taxonomic assignments for 16S ASVs. The top BLAST ID was used to assign identity for further filtering and analysis.

| ASV | BEExact ID | Silva ID | BLAST ID |
|---|---|---|---|
| 85 | Alphaproteobacteria | Mitochondria | Mitochondria |
| 89 | Pseudomonadota | Mitochondria | Mitochondria |
| 101 | Bacteria | Mitochondria | Mitochondria |
| 111 | Pseudomonadota | Mitochondria | Mitochondria |
| 130 | Alphaproteobacteria | Mitochondria | Mitochondria |
| 175 | Pseudomonadota | Mitochondria | Mitochondria |
| 249 | Cyanobacteria | Chloroplast | Chloroplast |
| 349 | Cyanobacteria | Chloroplast | Chloroplast |
| 504 | Mitochondria | Chloroplast | Chloroplast |
| 534 | Bacteria | Mitochondria | Mitochondria |
| 632 | Bacteria | Mitochondria | Mitochondria |
| 865 | Bacteria | Archaea | Archaea |
| 956 | Bacteria | NA | Fungi |
| 970 | Bacteria | Mitochondria | Mitochondria |
| 972 | Cyanobacteria | Chloroplast | Bacteria |
| 975 | Bacteria | Archaea | Archaea |
| 978 | Bacteria | Mitochondria | *No Match* |
| 987 | Bacteria | Archaea | Archaea |
| 1026 | Bacteria | NA | Fungi |
| 1029 | Bacteria | Chloroplast | Algae |

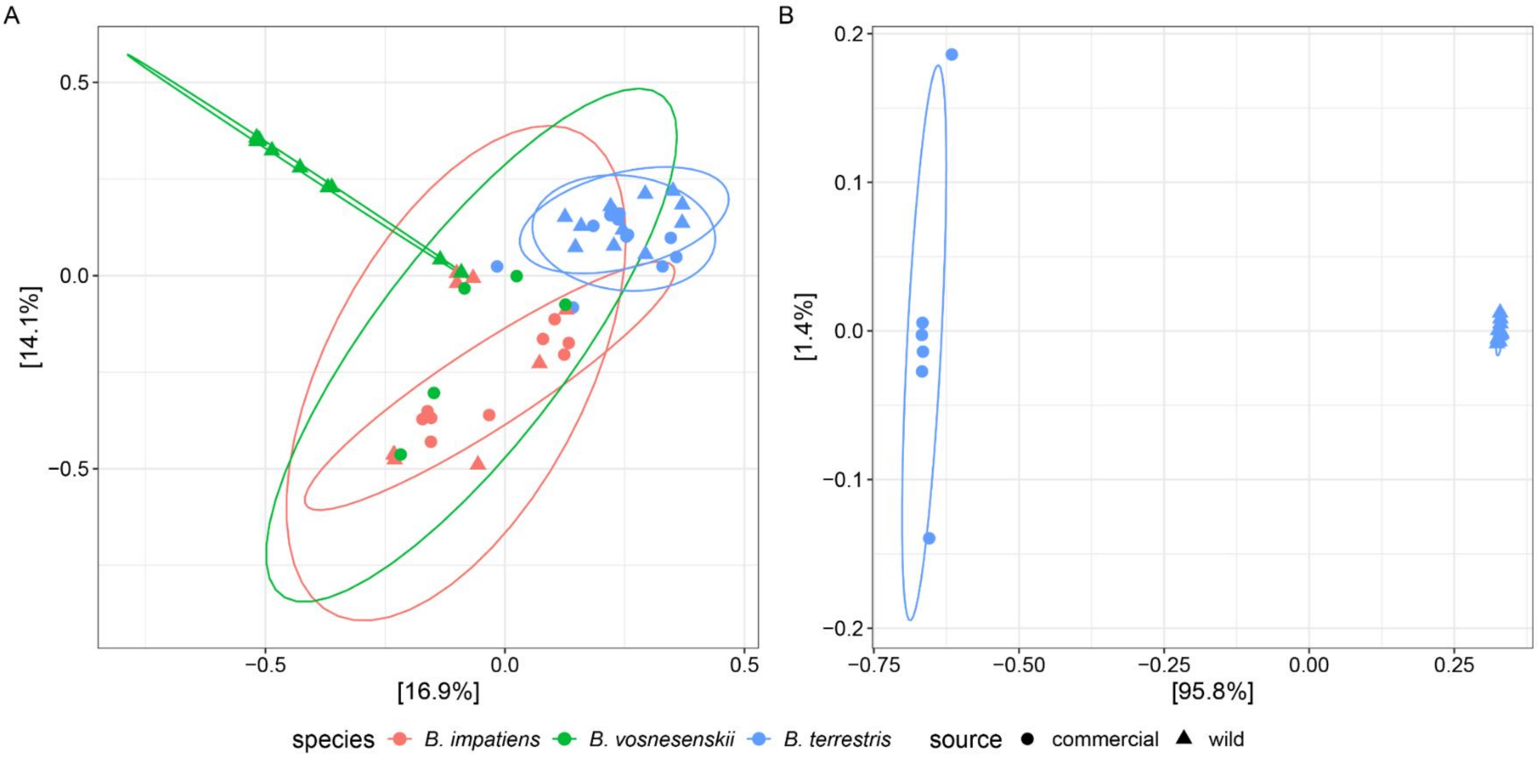


Figure S1. PCoA plots of bacterial communities of three species of bumble bees (A) and fungal communities of *B. terrestris* (B) from wild and commercial sources. The composition of bacterial communities differed by bee species ($F_{2,52}$ = 9.11, $R^2$ = 0.21, p = 0.001), source ($F_{1,52}$ = 5.61, $R^2$ = 0.066, p = 0.001), and their interaction ($F_{2,52}$ = 4.93, $R^2$ = 0.11, p = 0.001), while variance was unaffected by species or source. For fungal communities, composition was impacted by bee source ($F_{1,16}$ = 357, $R^2$ = 0.96, p = 0.001), while variance was unaffected by source.

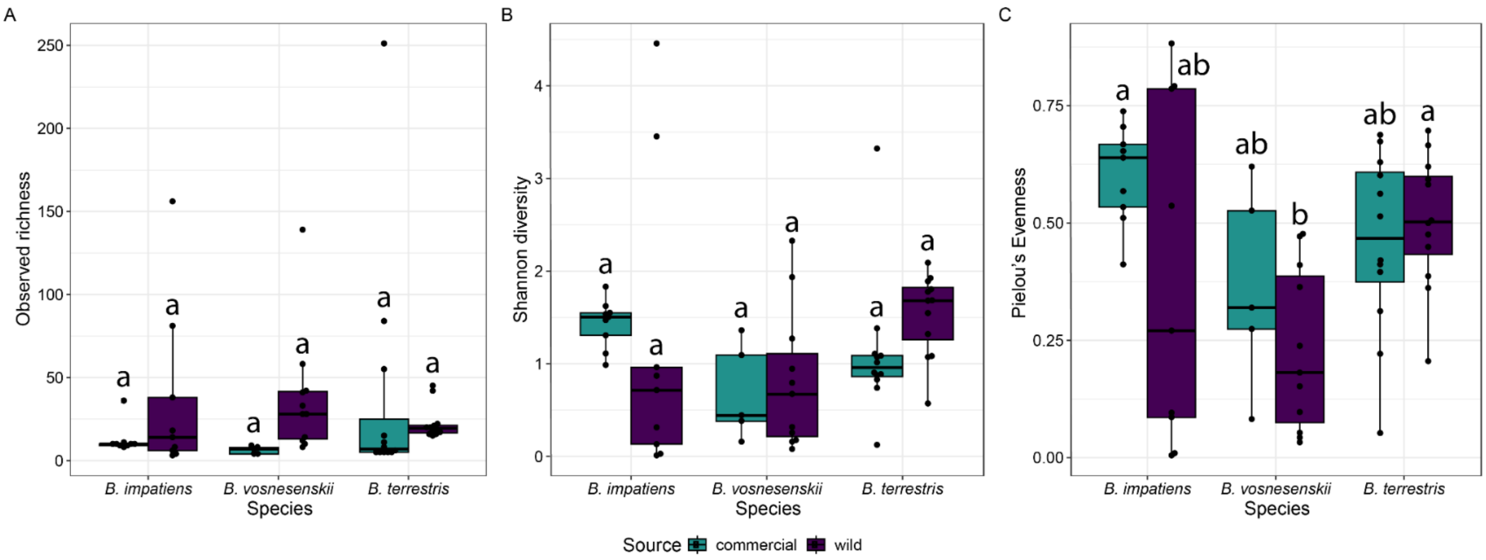


Figure S2. Observed richness (A), Shannon diversity (B), and Pielou's evenness (C) of total bacterial communities associated with three species of bumble bees from commercial and wild sources. Observed ASV richness was unaffected by bee species ($F_{2,52}$ = 0.095, p = 0.91), source ($F_{1,52}$ = 0.603, p = 0.44), or their interaction ($F_{2,52}$ = 1.74, p = 0.19). Overall Shannon diversity was also unaffected by source ($F_{1,52}$ = 0.40, p = 0.53) or the species * source interaction ($F_{2,52}$ = 0.76, p = 0.47), and was only marginally impacted by bee species ($F_{2,52}$ = 2.50, p = 0.092). Community evenness was lower in *B. vosnesenskii* samples than in *B. impatiens* or *B. terrestris* ($F_{2,52}$ = 6.14, p = 0.004), but was unaffected by bee source ($F_{1,52}$ = 2.25, p = 0.14), or the species * source interaction ($F_{2,52}$ = 0.2.2, p = 0.12). Different letters represent statistically different values as determined by estimated marginal means (α = 0.05).

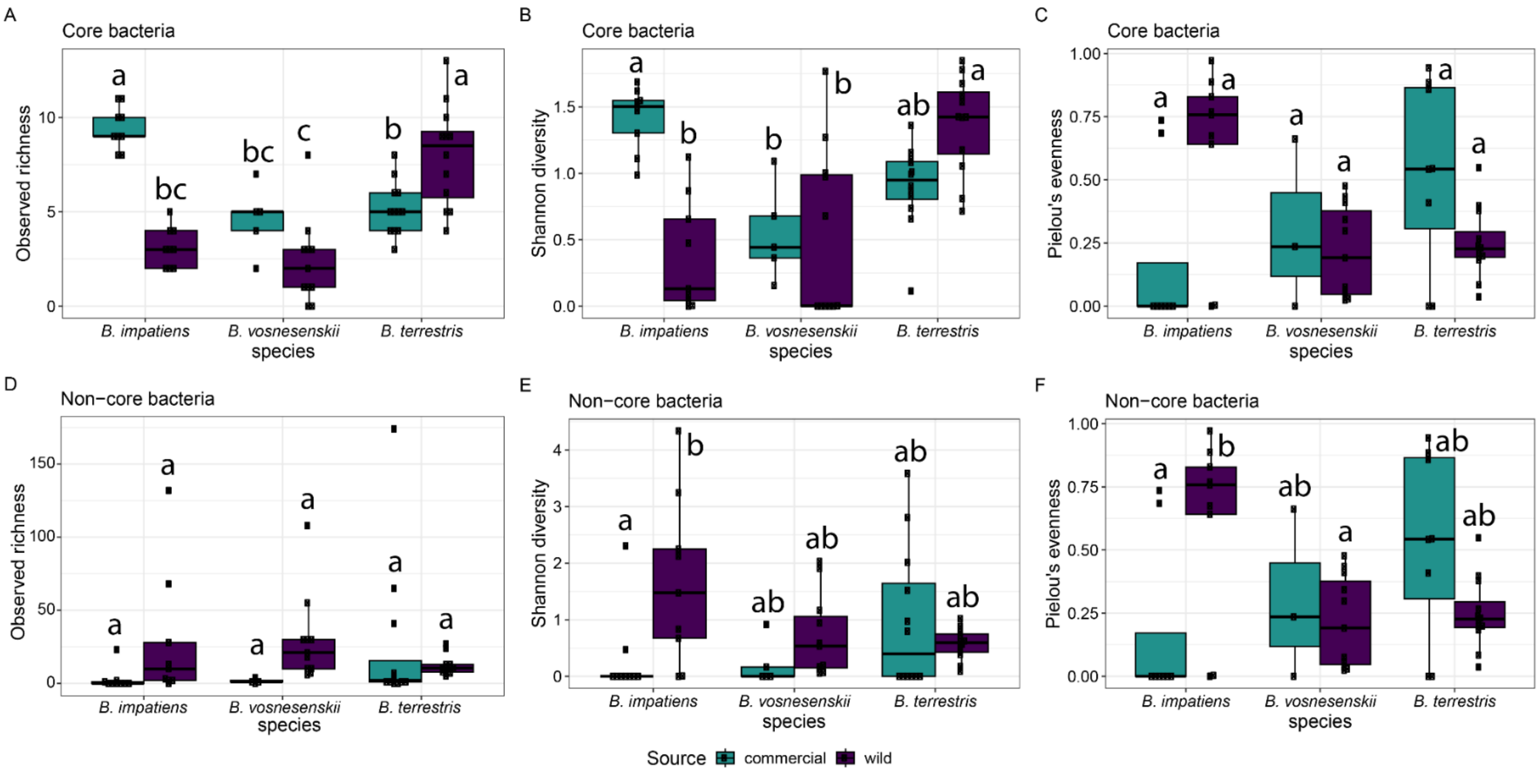


Figure S3. Observed richness, Shannon diversity, and Pielou's evenness of core (A, B, C) and non-core (D, E, F) bacterial taxa associated with three species of bumble bees from commercial and wild sources. For core taxa, observed richness was impacted by species ($F_{2,52} = 18.7$, $p < 0.001$), source ($F_{1,52} = 7.4$, $p < 0.01$), and their interaction ($F_{2,52} = 30.6$, $p < 0.001$). Shannon diversity of core bacteria was affected by species ($F_{2,52} = 10.2$, $p < 0.001$) and the source * species interaction ($F_{2,52} = 16.9$, $p < 0.001$), while Pielou's evenness was unaffected by all factors (Species - $F_{2,49} = 1.12$, $p = 0.33$; Source - $F_{1,49} = 0.04$, $p = 0.84$; Interaction - $F_{2,49} = 2.8$, $p = 0.071$). For non-core taxa, Shannon diversity ($F_{2,52} = 4.5$, $p = 0.02$) and Pielou's evenness ($F_{2,45} = 4.47$, $p = 0.002$) were affected by the source * species interaction, while observed richness was unaffected by any tested factors ($p > 0.05$ for all comparisons). Different letters represent statistically different values as determined by estimated marginal means ($\alpha = 0.05$).

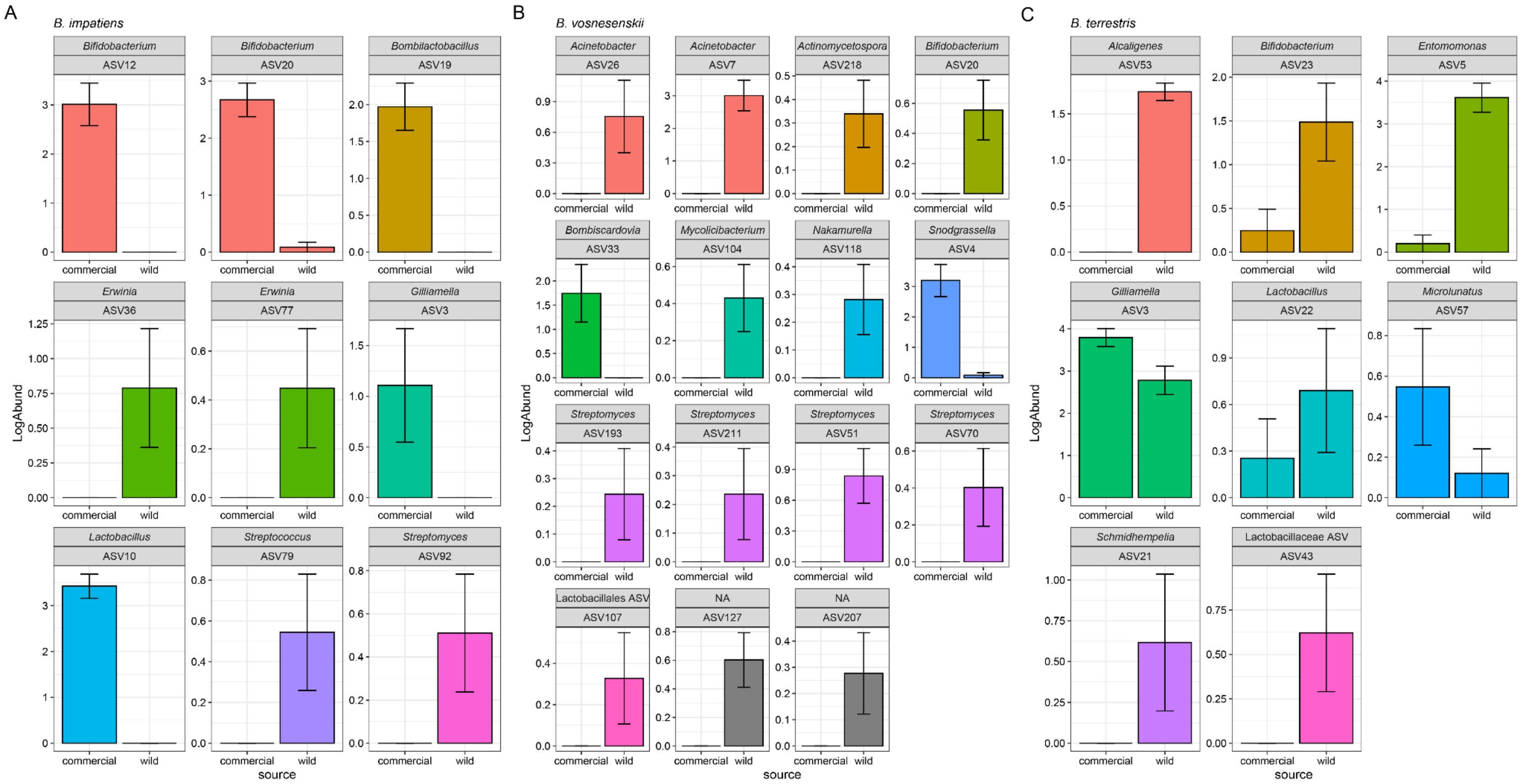


Figure S4. Differently abundant bacterial taxa between commercially-reared and wild-caught bumble bees belonging to three species as determined by differential gene expression analysis using DESeq2 ($\alpha = 0.05$).

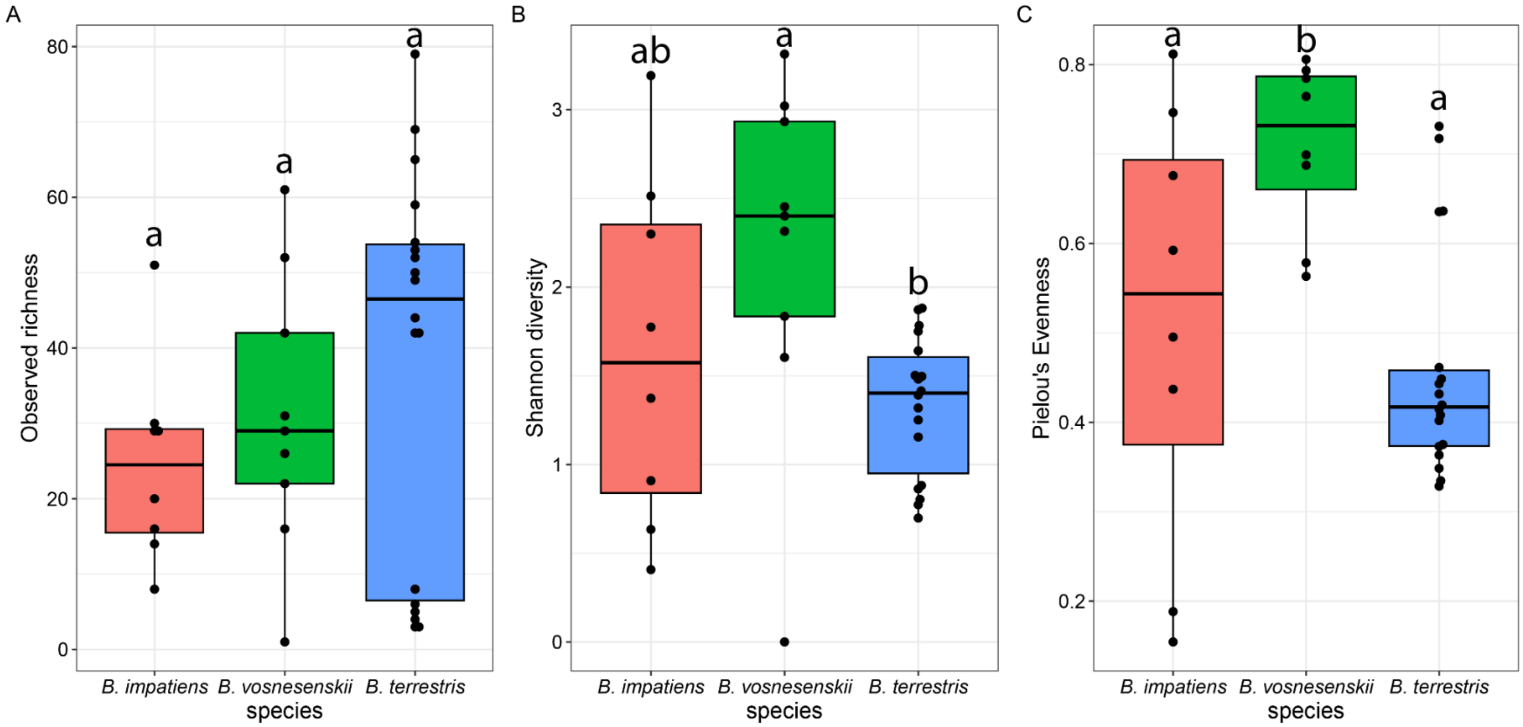


Figure S5. Diversity of fungal gut communities associated with three species of bumble bees. Species had no impact on observed ASV richness ($F_{2,32} = 1.11$, $p = 0.34$), but did impact Shannon diversity ($F_{2,32} = 4.28$, $p = 0.023$) and community evenness ($F_{2,31} = 7.1$, $p = 0.003$). Different letters represent statistically different values as determined by estimated marginal means ($\alpha = 0.05$).

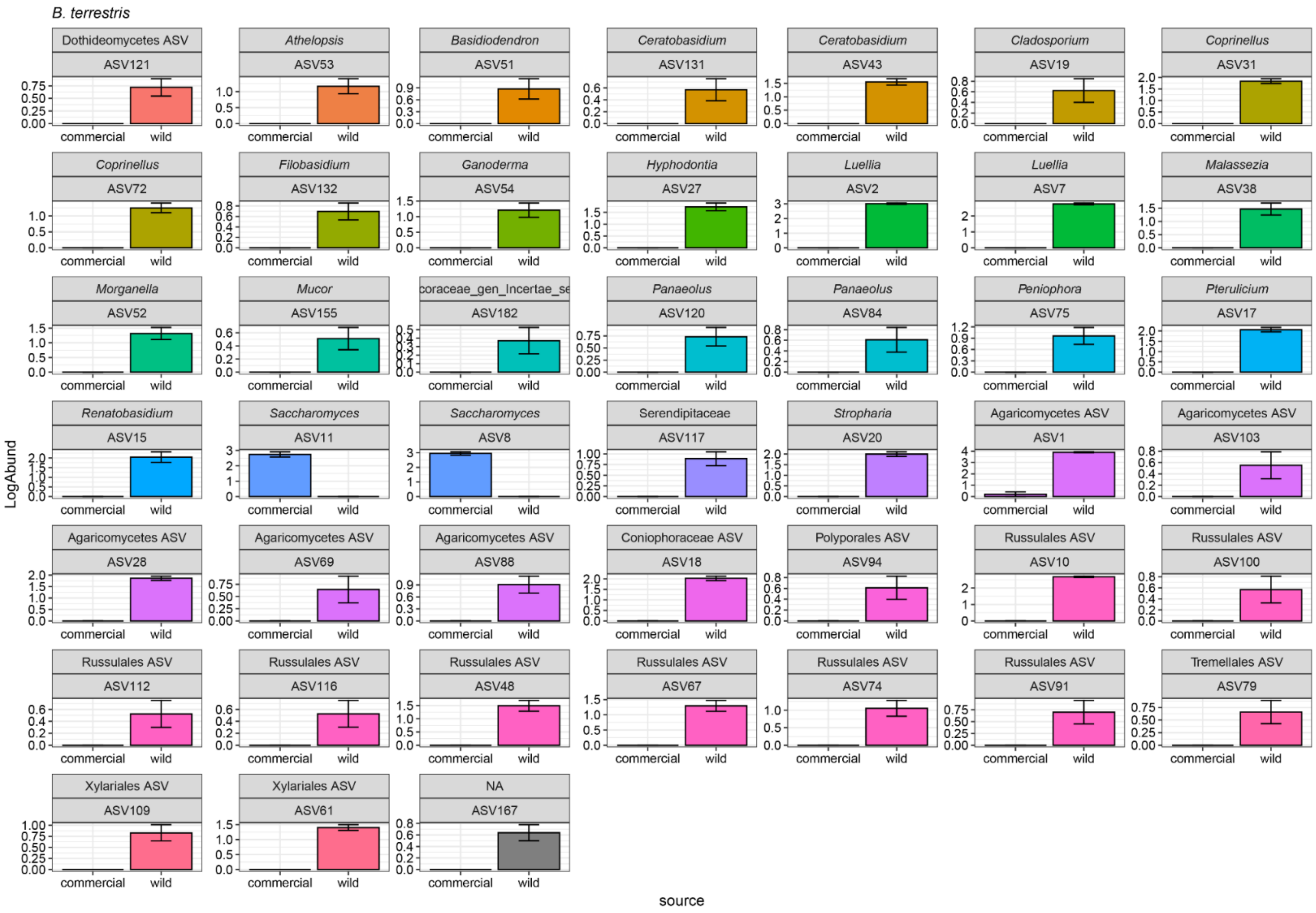


Figure S6. Differently abundant fungal taxa between commercially-reared and wild-caught workers of *B. terrestris* as determined by differential gene expression analysis using DESeq2 (α = 0.05).

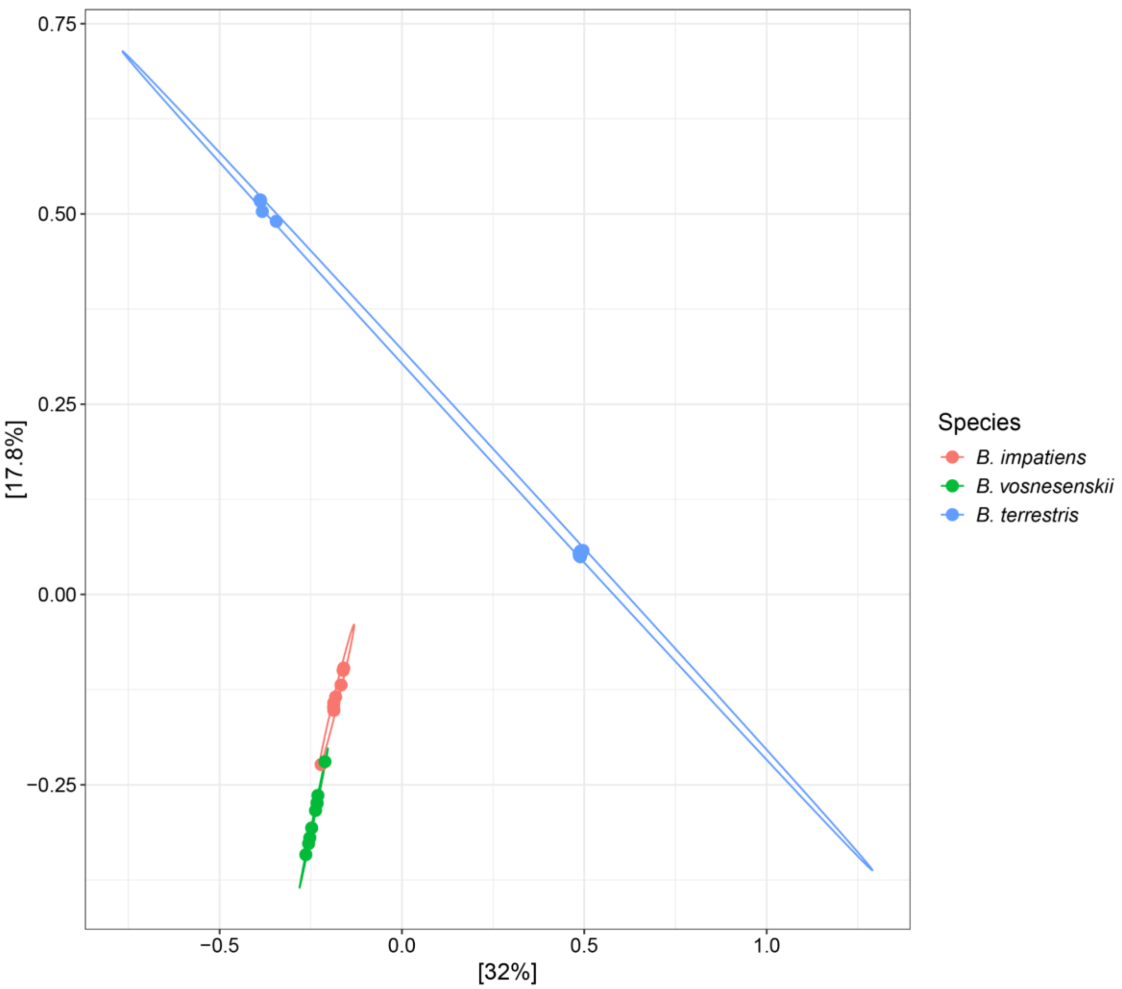


Figure S7. PCoA plots of fungal communities of three species of bumble bees. Community composition varied between bee species ($F_{2,32}$ = 6.94, $R^2$ = 0.30, p = 0.001), while variance was similar across species ($F_{2,32}$ = 2.12, p = 0.13)
